# Brachyury cooperates with Wnt/β-Catenin signalling to elicit Primitive Streak like behaviour in differentiating mouse ES cells

**DOI:** 10.1101/003871

**Authors:** David A. Turner, Pau Rué, Jonathan P. Mackenzie, Eleanor Davies, Alfonso Martinez Arias

## Abstract

The formation of the Primitive Streak is the first visible sign of gastrulation, the process by which the three germ layers are formed from a single epithelium during early development. Embryonic Stem Cells (ESCs) provide a good system to understand the molecular and cellular events associated with these processes. Previous work, both in embryos and in culture, has shown how converging signals from both Nodal/TGFβR and Wnt/β-Catenin signalling pathways specify cells to adopt a Primitive Streak like fate and direct them to undertake an epithelial to mesenchymal transition (EMT). However, many of these approaches have relied on genetic analyses without taking into account the temporal progression of events within single cells. In addition, it is still unclear as to what extent events in the embryo are able to be reproduced in culture. Here, we combine flow-cytometry and a quantitative live single-cell imaging approach to demonstrate how the controlled differentiation of mouse ESCs (mESCs) towards a Primitive Streak fate in culture results in cells displaying many of the characteristics observed during early mouse development including transient Brachyury expression, EMT and increased motility. We also find that the EMT initiates the process, and this is both fuelled and terminated by the action of Bra, whose expression is dependent on the EMT and ß-Catenin activity. As a consequence of our analysis, we propose that a major output of Brachyury expression is in controlling the velocity of the cells that are transiting out of the Primitive Streak.

## Introduction

The development of an organism is the result of the proliferation, concomitant phenotypic diversification and spatial organisation of cells in the context of spatiotemporally controlled patterns of gene expression(Arias and Stewart, 2002; Gilbert, 2013; Wolpert and Tickle, 2010). The use of genetics to interrogate these processes has revealed that they are underpinned by the temporal iteration of coordinated interactions between signal transduction and transcription factor networks. Establishing the relationship between these molecular events and the emergence of cellular diversity is an essential step towards understanding the relationship between programs of gene activity and the process of morphogenesis that shapes cells into tissues and organs.

Gastrulation is one of the earliest events where it is possible to observe a convergence of fate specification and morphogenetic processes in embryos (Wolpert and Tickle, 2010). It occurs in all metazoan and encompasses a choreography of cell movements that transforms a group of seemingly identical epithelial cells, with species specific geometry, into the outline of an organism exhibiting an overt anterior-posterior organisation and three germ layers (ectoderm, mesoderm, endoderm) (Ramkumar and Anderson, 2011; Solnica-Krezel and Sepich, 2012; Tam and Gad, 2004). In chordates, gastrulation is led by a dynamic population of cells that gives rise to the mesoderm and the endoderm, defines and patterns the neuroectoderm and delineates the plane of bilateral symmetry (Nowotschin and Hadjantonakis, 2010; Tam and Gad, 2004). In mammalian embryos this population, called the Primitive Streak, is associated with the expression of the T-box transcription factor Brachyury (Bra) (Beddington et al., 1992; Fehling et al., 2003; Ramkumar and Anderson, 2011; Rivera-Pérez and Magnuson, 2005; Wilkinson et al., 1990). In the mouse, Bra is first expressed shortly after implantation in a group of cells at the boundary between the prospective embryonic and extraembryonic tissues. At stage E6.5, preceding the onset of gastrulation movements, Bra expression becomes restricted to the proximal posterior region of the embryo (Rivera-Pérez and Magnuson, 2005), at the position where under the influence of Nodal and Wnt signalling, the Primitive Streak is initiated as a dynamic structure that will progress towards the distal end of the epiblast, ploughing an anteroposterior axis (Arnold and Robertson, 2009). At the cellular level, gastrulation involves a sequence of highly organized Epithelial Mesenchymal Transition (EMT) movements that propagate through the tissue in a manner that resembles a travelling wave (Lim and Thiery, 2012; Thiery and Sleeman, 2006; Williams et al., 2011). The first cells undergoing EMT invaginate and move towards the anterior contralateral side of the embryo. This movement accompanies the distal/anterior spread of the streak and thus, by the end of gastrulation, two thirds of the epiblast have been wrapped by the cells that have undergone gastrulation. This choreography of cell movements is characterized by the expression of a number of transcription factors at the leading edge of the EMT, in particular Brachyury (Wilkinson et al., 1990), Eomes (Russ et al., 2000) and Mixl1 (Robb et al., 2000).

Just anterior to the Primitive Streak there is a structure, the organizer, which does not undergo an EMT, shows homology with the Spemann organizer and will become the Node when the streak reaches the distalmost anterior region of the epiblast at about E7.5 (Beddington, 1994). The organizer and the Node express Bra and, after E7.5 the Node regresses towards the posterior pole of the embryo, leaving the notochord in its wake, the Node and the notochord, continue to express Bra. After this time, Bra expression becomes restricted to a region in the tail that will undergo caudal extension to generate the caudal spinal cord and somatic mesoderm, (Beddington et al., 1992; Cambray and Wilson, 2007; J. C. Smith, 2004; Tam and Gad, 2004; Wilson et al., 1993). Genetic analysis has shown that in embryos, Bra is required for both movement of the cells through the Primitive Streak during axial extension and their specification into mesoderm and notochord posterior to somite 7 (Gluecksohn-Schoenheimer, 1938; B. L. Martin and Kimelman, 2008; Wilson et al., 1995).

The localization of Bra expression to the proximal posterior region of the epiblast is a useful reference for the onset of gastrulation and genetic analysis has identified a requirement for BMP, Wnt/β-Catenin, and Nodal for this event (Tam and Loebel, 2007). These studies have also shown that BMP sets up the expression of Nodal and Wnt3a, which, from the visceral endoderm, are likely to be the direct activators of Bra expression. How individual cells integrate Nodal and Wnt signalling with the expression of Bra to promote the directional EMT and how these events are coordinated across the cell population that is defined as the Primitive Streak is not known. While there are some reports of the gastrulation movements in mouse embryos with a cellular resolution (Ichikawa et al., 2013; Williams et al., 2011), these are descriptive and the visualization systems do not lend themselves to experimental perturbations.

Embryonic Stem Cells (ESCs), clonal populations of cells from the pre-implantation blastocysts (M. J. Evans and Kaufman, 1981; G. R. Martin, 1981), can be differentiated in culture into Bra expressing cells that exhibit gene expression profiles characteristic of the Primitive Streak (Fehling et al., 2003; Gadue et al., 2006; Hansson et al., 2009; Keller, 2005; Kubo et al., 2004; Ying et al., 2003a; Ying and A. G. Smith, 2003; Ying et al., 2003b; 2008). In addition to gene expression analysis, ESCs also offer the opportunity to quantify proteins at the level of single-cells through live imaging. For these reasons ESCs could provide a useful model to understand the link between signals and morphogenesis in the context of the onset of gastrulation. However, in order to do this it is important to show that, in addition to patterns of gene expression, the differentiating ES cells share other features with the cells in the Primitive Streak, in particular EMT and its relationship to Bra expression.

Here we have used a combination of live cell imaging, immunocytochemistry and chemical genetics to analyse the onset and consequences of Bra expression in mESCs at the level of single cells. We observe that ES cells grown on gelatin and in the presence of Activin (Act) and Chiron (Chi, an agonist of Wnt/β-Catenin signalling), undergo an EMT associated with the expression of Bra and that the EMT itself contributes to Bra expression. We are able to separate the inputs of β-Catenin and Act in the onset of Bra expression and show that while β-Catenin is required for the up-regulation of Bra, Act is required for the velocity, and thereby the distance cells travel. We discuss our findings in the context of the emergence of the Primitive Streak during gastrulation and suggest that differentiation of ES cells into Bra expressing cells in culture provides a valuable system to study the mechanisms that specify the emergence of the Primitive Streak.

## Materials and Methods

### Routine Cell Culture and Primitive Streak Differentiation

E14Tg2A, Bra::GFP (Fehling et al., 2003), Bra^-/-^ (Beddington et al., 1992), TLC2 (Ferrer-Vaquer et al., 2010), and Nanog^-/-^ (Chambers et al., 2007) mESCs were cultured on gelatin in serum and LIF (ESLIF). β-Catenin^-/-^ and β-Catenin^ΔC^ mECS (Wray et al., 2011) were cultured in 2iLIF (Ying et al., 2008) on fibronectin. For differentiation assays, cells were plated in ESLIF (6×10^3^) and 24 hours later medium replaced with N2B27 (Ying and A. G. Smith, 2003) (NDiff 227, StemCells Inc., UK) supplemented with LIF & BMP. Cells were differentiated 24 hours later in N2B27 with combinations of ActivinA (Act; 100 ng/ml), CHIR99021 (Chi; 3 μM) or dual stimulation (Act/Chi). Differentiation of β-Catenin^-/-^ and β-Catenin^ΔC^ lines omitted the LB stage. Small molecule inhibitors used were 1 μM XAV939 (Huang et al., 2009); 1 μM IWP3 (Chen et al., 2009); CyclosporineA (CsA (Li et al., 2011); 5 μM), SB431542 (SB43 (Inman et al., 2002); 10 μM) and Dorsomorphin (DM (P. B. Yu et al., 2007); 0.2 μM). If cells were not being passaged, they were fed daily by replacing half the medium in the tissue culture flask with fresh growth medium.

### FACS analysis

GFP and RFP was assessed using a Fortessa flow cytometer (BD Biosciences). Analysis of data from single, live cells (DAPI-negative) was conducted using Flowjo software (Tree Star, Inc.). The GFP-positive populations from FACS data were analysed using a one-way ANOVA with Tukey's adjustment comparing the time-matched DMSO control and treatment; significance was set at *p* < 0.05.

### Quantitative Image Analysis (QIA) and Confocal Microscopy

Immunofluorescence and image analysis carried out as described previously (Descalzo et al., 2012) in 8-well (Ibidi), plastic tissue-culture dishes. Primary antibodies are described in the supplemental material. Samples imaged using and LSM700 on a Zeiss Axiovert 200M with a 40x EC Plan-NeoFluar 1.3 NA DIC oil-immersion objective. Hoechst, Alexa-488, -568 and - 633/647 were sequentially excited with a 405, 488, 555 and 639 nm diode lasers respectively. Data capture was carried out using Zen2010 v6 (Zeiss) and image analysis performed using Fiji (Schindelin et al., 2012).

### Widefield, live-cell Microscopy and analysis

For live imaging, cells were imaged by widefield microscopy in a humidified CO_2_ incubator (37°C, 5% CO_2_) every 10 minutes for up to 96 hours using a 20x LD Plan-Neofluar 0.4 NA Ph2 objective with correction collar set to image through tissue-culture plastic dishes. An LED, white-light system (Laser2000) excited fluorescent proteins. Emitted light was recorded using an AxioCam MRm and recorded with Axiovision (2010) release 4.8.2. Analysis performed using Fiji (Schindelin et al., 2012) and associated plugins: MTrackJ (Meijering et al., 2012) or Circadian Gene Expression (CGE) (Dibner et al., 2009; Sage et al., 2010). When MTrackJ was used, the bright-field channel from the live cell imaging was used to manually identify cell nuclei, the centre of which was used as a seed point for the position of the cells. Single cells were then tracked until the cell divided, became obscured by other cells or left the field of view. Supplementary Table S1 summarises the tracking data collected. To measure the level of reporter expression, a small circular region of interest of radius 10μm was automatically generated around the centre of each cell and the average pixel intensity from the fluorescence channel was recorded. Movies not included in the supplementary data are available upon request.

### Quantitative RTPCR analysis

Cells were harvested at the relevant time-points, RNA extracted in TRIzol (Life Technologies, UK) and purified using the Qiagen RNeasy Mini Kit (Qiagen). Complementary DNA (cDNA) synthesis was performed with 1 µg of total RNA extract using the SuperScript III First-Strand Synthesis System (Life Technologies). RNA was digested afterwards with RNaseH. PCR reactions were run in triplicates using 12.5 µl QuantiFAST SYBR Green (Qiagen) Master Mix, 3 µl cDNA, 0.5 µl primer mix (50 µM) and 9 µl dH_2_O. Pipetting was performed automatically by QIAgility (Qiagen). Initial sample concentrations were estimated using an in-house adapted MAK2 method (Boggy and Woolf, 2010); technical repeats were averaged and normalised to GAPDH levels. Standard errors were propagated accordingly. Primer pairs are as follows: Brachyury forward CTGGGAGCTCAGTTCTTTCG, reverse GTCCACGAGGCTATGAGGAG; GAPDH forward AACTTTGGCATTGTGGAAGG, reverse GGATGCAGGGATGATGTTCT. Cycling conditions: initial denaturation at 95°C for 5 minutes, followed by 45 cycles of denaturation at 95°C for 10 seconds and combined annealing and extension at 60°C for 30 seconds. Melt curve generated between 60 and 95°C, holding for 5 seconds for each temperature step.

## Results

### The Onset of Brachyury Expression in culture

To probe the connection between the differentiation of mouse embryonic stem cells (mESCs) and the onset of Bra expression in the embryo, we used a mESC line bearing an insertion of GFP into the Bra locus (Bra::GFP) that has been shown to display, upon differentiation, characteristics associated with the Primitive Streak during development (Fehling et al., 2003; Gadue et al., 2006). We confirmed that this line faithfully reports the onset of Bra expression in culture by following, through FACS analysis, Bra::GFP expression in a variety of differentiation conditions (Fig. 1A) and comparing its expression profile to that of endogenous Bra protein and mRNA (Fig. 1B,C). When Bra::GFP-expressing cells are grown in neural differentiation conditions, there is no GFP expression (Fig. S1) but growth in the presence of ActivinA (Act) and an inhibitor of GSK3, CHIR99021 (Chi), that mimics Wnt/ß-Catenin signalling) leads to a wave of Bra expression (Fig. 1Ai) that parallels Bra mRNA and protein (Fig. 1B, C). As this cell-line is heterozygous for Bra, it is likely to exhibit an haploinsuficient phenotype associated with the Bra locus (Dobrovolskaia-Zavadskaia, 1927; Rashbass et al., 1991; Wilson et al., 1993), but this should not interfere with the initiation of expression, which is the object of our study.

**Figure 1:**
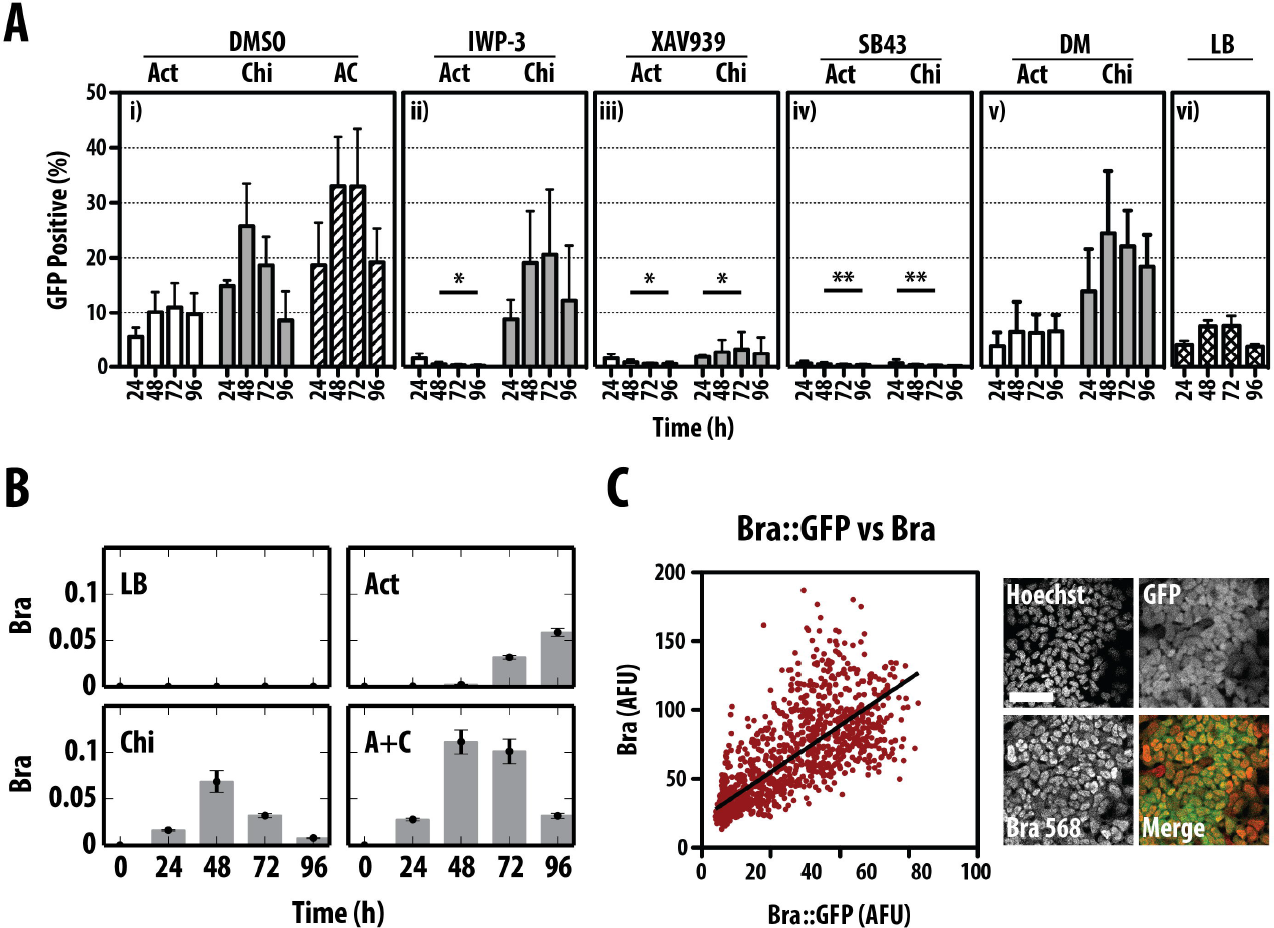
Induction of Bra expression by Activin (Act) and CHIR99021 (Chi) during the differentiation of mESCs. (A) Bra::GFP mESCs were treated with Act, Chi or Act/Chi and DMSO (A-i), Act and Chi with the Porcupine inhibitor IWP3, which inhibits the secretion of Wnt proteins (A-ii), with the Tankyrase inhibitor XAV939, which reduces active ß-Catenin (A-iii), the Nodal/Act receptor inhibitor SB43 (A-iv) or the BMP inhibitor Dorsomorphin (A-v). A control for long term pluripotency growth, LIF and BMP (LB) is included (A-vi). Notice that the robust expression of Bra induced by Act and Chi, is suppressed by inhibition of Wnt/ß-Catenin or Nodal/Act signalling but not by BMP inhibition. GFP-positive cells measured daily by FACS (± S.D. from at least 3 replicates). (B) E14-Tg2A mESCs were grown in LB, Act, Chi or Act/Chi for the indicated durations prior to RNA extraction and RT-qPCR analysis for the indicated genes. The average expression level of the RT-qPCR replicates relative to that of GAPDH are shown for a representative experimental replicate. Error bars indicate absolute error of the normalized mean. Notice the transient expression of Bra in Chi and dual Act and Chi compared with the delayed expression in Act (C) Quantification of Bra::GFP v Bra expression by immunostaining indicates a high correlation (Pearson coefficient of 0.773); scale bar denotes 50 μm. Single and double asterisk denotes *p* < 0.05 & *p*<0.01 respectively versus DMSO.

Individual treatment of the Bra::GFP mESCs with either Act (100 ng/ml) or Chi (3 μM) resulted in a transient rise in the proportion of cells expressing GFP (Fig. 1Ai). In the case of Act alone we observe a peak of ∼10% at 72 hours (Fig. 1Ai), whereas treatment with Chi anticipates this peak with 25% of the total population expressing Bra::GFP at 48 hours (Fig 1Ai). The combination of Act and Chi (Act/Chi) results in an increase in the proportion of cells expressing Bra::GFP (∼33% GFP positive at 48-72h) which is sustained relative to Chi alone (Fig. 1Ai). These observations, extend previous ones (Gadue et al., 2006; Kubo et al., 2004) and suggest that both Act and Wnt/β-Catenin signalling can influence the progression of pluripotent cells towards a state characterized by the expression of Bra.

To understand the individual contributions of Act and Wnt/β-Catenin signalling to the expression of Bra::GFP, specific inhibitors targeting each pathway were added to the cell culture medium (Fig. 1Aii-v). Inhibition of β-Catenin signalling during Act treatment (Fig. 1Aii,iii) resulted in complete ablation of Bra::GFP induction compared with the DMSO control (*p-value* < 0.05). A similar effect was observed upon inhibition of the Act signalling by SB43 (an inhibitor of Act/Nodal signalling) in the presence of Act or Chi (Fig. 1Aiv).

Taken together, these results are in agreement with previous observations (Gadue et al., 2006) that the onset of Bra::GFP by Act ligands not only requires active β-Catenin signalling, but that an active Act pathway is absolutely required for β-Catenin-mediated Bra induction.

### An EMT associated with mesendodermal differentiation in ES cells

In the embryo, the onset of Bra expression is associated with spatiotemporally organized cellular movements that configure the process of gastrulation (Lim and Thiery, 2012; Nakaya et al., 2008) In order to assess whether this behaviour was also observed in ESCs in these conditions, we analysed the cellular activities associated with Bra expression in the adherent cultures by filming the behaviour of the cells in the presence of Act/Chi (Fig. 2A), a condition that promotes the maximal Bra expression response (Fig. 1Ai). We observe that after 24 hours, the colonies characteristic of the pluripotent state loosen up and the differentiating cells form an epithelial monolayer from which they adopt a mesenchymal-like morphology and become motile (Fig. 2A). This state is probably akin to the postimplantation epiblast and it is in this state that it is possible to observe the onset of Bra expression. This sequence of events is reminiscent of Epithelial to Mesenchymal Transitions (EMT), which characterize cells during the process of gastulation. To investigate this further we probed for phenotypic landmarks of the EMT process in the differentiating cells (Fig. 2B-D).

**Figure 2:**
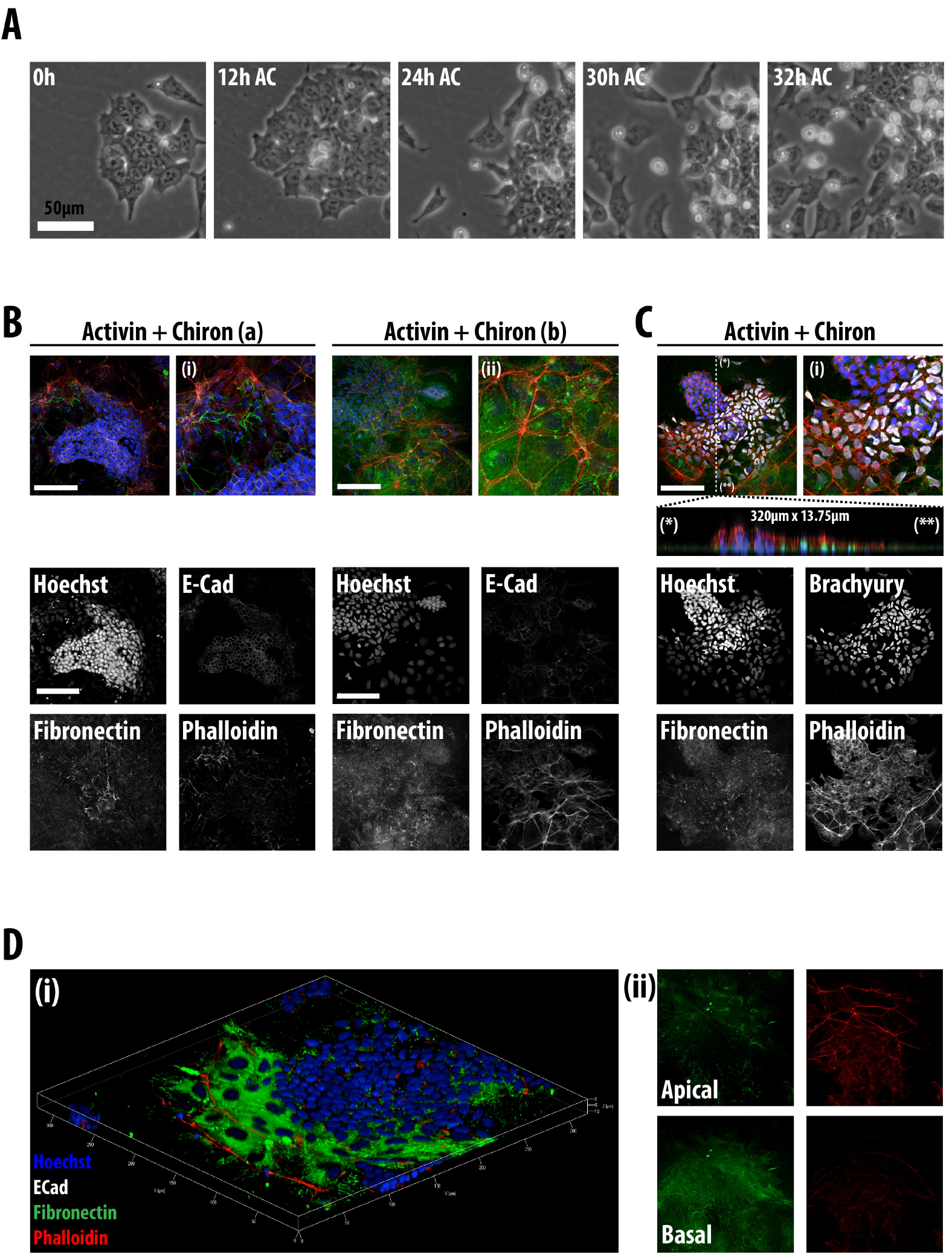
An EMT is associated with mesendodermal differentiation of E14-Tg2A. (A) Phase-contrast still images from live cell imaging of wild-type mESCs treated with Act/Chi. At the start of differentiation, cells begin to loosen up within the colonies and as time progresses adopt a more motile morphology and become migratory. (Ba,b & C) Differentiating mESCs were treated with Act/Chi and stained for E-Cadherin (B) or Brachyury (C) together with Fibronectin and Phalloidin. Panels (a) and (b) in (B) show two separate colonies and their corresponding magnified regions (i and ii) in different phases of an EMT. The decrease in E-Cadherin correlates with the appearance of filopodia and the laying down of Fibronectin basally. This is clear at the edge of the colony, where these changes are associated with the expression of Bra (C). This is also illustrated when a slice through the colony (indicated by * and **) in the z-dimension shows the different phases of the EMT (middle, horizontal panel in C). (D) A three-dimensional rendering of E14-Tg2A mESCs in the presence of Act/Chi, stained for E-Cadherin (white), Fibronectin (green) and Phalloidin (red). Cells emerging from the colony secrete high levels of Fibronectin and an altered F-actin architecture. Part (ii) reveals the location of Fibronectin on the basal surface of the colony. Scale bar corresponds to 50 µm in (A) and 100 μm in (B-D). Hoechst is used to stain the nuclei in all images.

Pluripotent cells are characterized by high levels of E-Cadherin (Chou et al., 2008; Malaguti et al., 2013; Soncin et al., 2009) even though they lack an epithelial structure and when grown on gelatin, exhibit low levels of Fibronectin basally (Fig. S2A,B). Analysis of fixed samples of cultures exposed to Act/Chi, shows that the onset of differentiation into mesendoderm is associated with a decrease in the levels of E-Cadherin concomitant with the onset of Bra expression (S2A). The expression of Bra tends to be localized to the edges of the disorganizing colonies and coincides with a rearrangement of the actin cytoskeleton and a boundary of expression of Fibronectin (see Fig 2C & S2C,D). As cells become motile, there is an increase in the amount of Fibronectin basally, as well as a progressive increase in the levels of F-Actin apically (Fig. 2B-D & Fig. S2C,D). A proportion of F-actin is found in the Lamellipodia and Filopodia at the edges of the colony in what appears to be moving cells (Fig. 2D & S2B).

These rearrangements of cellular elements are typical of EMT and are very similar to those of cells moving into the Primitive Streak in embryos during stage E.7.5 (See Fig. 2C and 3D in (Migeotte et al., 2011))(Nakaya and Sheng, 2008) suggesting that, under the influence of Act and Wnt signalling in culture, ES cells have activated a developmental programme very similar to that of cells undergoing gastrulation.

### Brachyury expression correlates with Nanog expression during ES cell differentiation

A hallmark of ES cell differentiation is the down-regulation in the expression of elements of the pluripotency network. To understand the relationship between this process and the onset of mesendodermal differentiation at the level of single cells, we used Quantitative Image Analysis (QIA) to monitor the expression of Bra in wild-type E14-Tg2A mESCs relative to that of Nanog, Oct4 and Sox2 following Act, Chi or Act/Chi treatment (Fig. 3 & Fig. S3).

**Figure 3:**
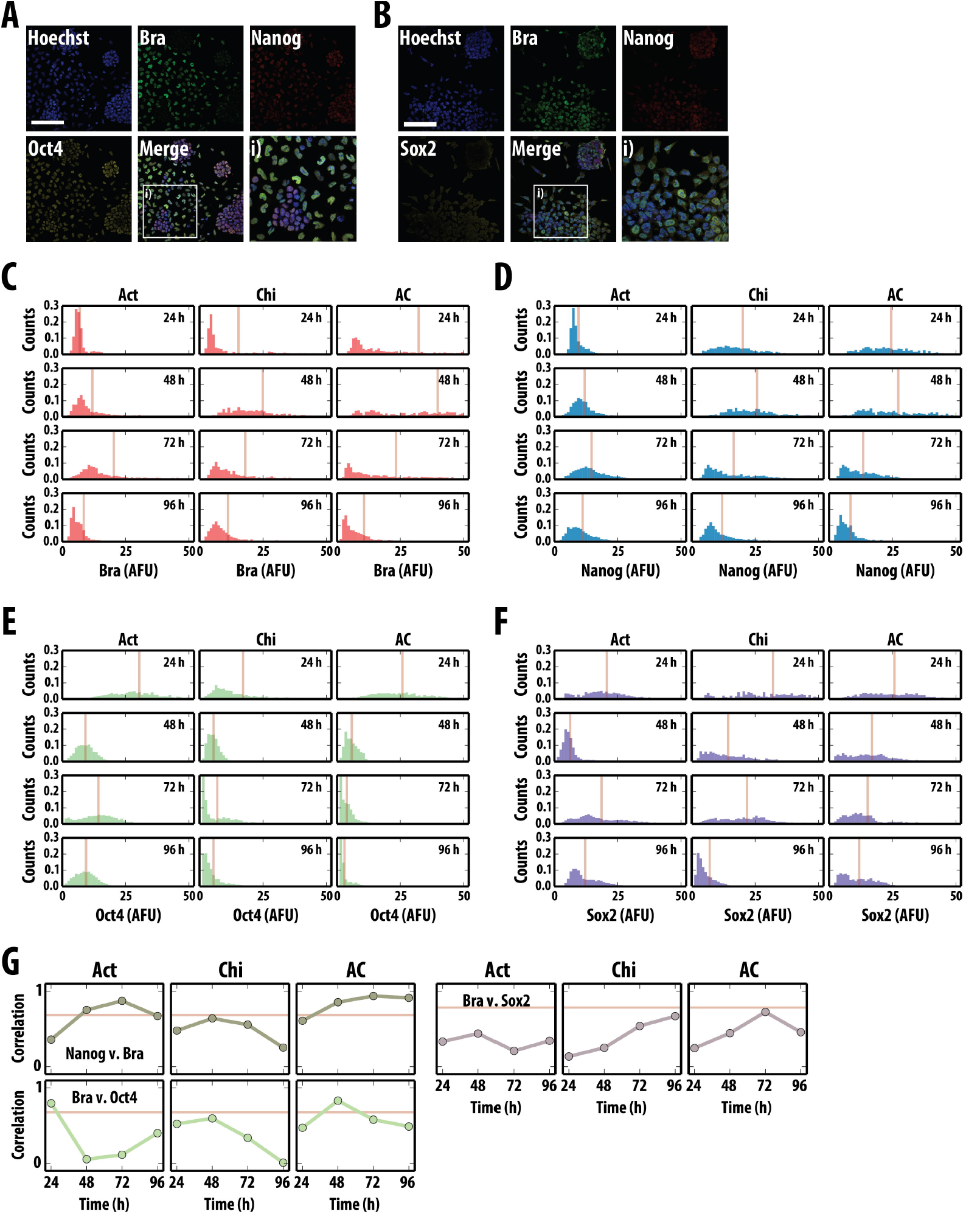
Quantitative Image Analysis (QIA) of Bra, Nanog and Oct4 or Sox2 following treatment with mesendodermal-inducing factors. (A,B) Confocal images of wild-type mESCs following 48h Act/Chi treatment stained for Brachyury (Bra; green) and Nanog (red) with either Oct4 (A) or Sox2 (B, yellow). Merged images shown with the corresponding magnified regions denoted by a white box (i). (C-F) Time evolution of the distributions of the expression of Brachyury (C), Nanog (D), Oct4 (E) and Sox2 (F) during differentiation. Wild-type E14-Tg2A mESCs treated with Act, Chi or Act/Chi for 24, 48, 72 and 96 hours were stained as described (A,B), nuclei segmented based on Hoechst staining and the average pixel intensity for each fluorescent channel (arbitrary fluorescence units: AFU) quantified. The intensities for Brachyury (C), Nanog (D), Oct4 (E) and Sox2 (F) are displayed as histograms for each time-point; the bisecting orange lines in each histogram corresponds to the mean fluorescence levels. (G) The Pearson Correlation Coefficient for the correlations between Bra and Nanog (top left), Bra and Oct4 (bottom left) and Bra and Sox2 (top left) for the different time points. The horizontal line represents the correlation for LIF BMP. Scale-bar represents 100μm; Hoechst is used to stain the nuclei.

Act treatment leads to the expression of Bra in a small proportion of the cells undergoing EMT whereas either Chi- or Act/Chi-induced differentiation resulted in a larger proportion of cells expressing Bra and, eventually, adopting a mesenchymal morphology. A small proportion of cells form tight balls around which the mesenchymal cells progressed (Fig. 3A,B). The cells in the tight balls expressed Nanog, Oct4 and Sox2 suggesting that within the colony centres, the cells remain pluripotent. Confocal images from these samples were then subject to QIA where the average pixel intensity in cell nuclei (identified and segmented by Hoechst uptake levels) was measured for each fluorescent channel (Fig. 3C-F).

Analysis of the time evolution of the distributions for the different proteins shows a wave of Bra expression in all conditions (Fig. 3C), though it is more pronounced and involves more cells in the case of Act/Chi. Chi alone also promotes the expression of Bra, but the levels and proportion of cells increase when it is supplemented with Act (Fig. 3C). In both cases, the increase in Bra levels is far from homogeneous within the population of cells, a fact that is compatible with imperfect coordination of its rapid and transient expression. We also noticed that while the levels of Oct4 and Sox2 decrease during the first two days of differentiation (Fig. 3E,F), in all cases Nanog expression undergoes a transient rise during days 2 and 3 (Fig. 3D). Analysis of pairwise protein levels in individual cells indicates an increase in the correlation between Nanog and Bra levels in Act and Act/Chi conditions which by day 2 go beyond the correlation levels in LIF BMP (Fig. 3G); a more transient and earlier increase in correlation levels can be observed between Oct4 and Bra in Act/Chi. In addition, strong correlations were also observed between Nanog and Oct 4 (Fig. S3), whereas Nanog and Sox2 became more correlated as time progressed (Fig. S3). These observations extend those of (Thomson et al., 2011) and suggest that the events in culture parallel the events in embryos where there is a transient rise in Nanog expression in the cells of the Primitive Streak at the start of gastrulation (Chambers et al., 2007; Hart et al., 2004; Osorno et al., 2012; Trott and Arias, 2013).

These results, together with those of the cell behaviours associated with the differentiation process, suggest that exposure of mESCs to Act/Chi triggers a developmental process that is very similar to that of the cells that undergo gastrulation in terms of gene expression (Gadue et al., 2006) as well as cell behaviour.

### The onset of Brachyury expression is tightly linked to an EMT and cell motility in adherent culture

Time-lapse imaging of Bra::GFP cells in different culture conditions (Fig. 4A & Fig. S4A) and subsequent manual tracking of single cells (Fig. 4B & Fig. S4B) allows us to simultaneously analyse the dynamics of Bra::GFP expression and the associated cell movement during the EMT, as well as the roles that Act and Wnt/β-Catenin signalling have individually in these processes at the level of single cells (Fig. 4 & Fig. S4; Movies M1,2).

**Figure 4:**
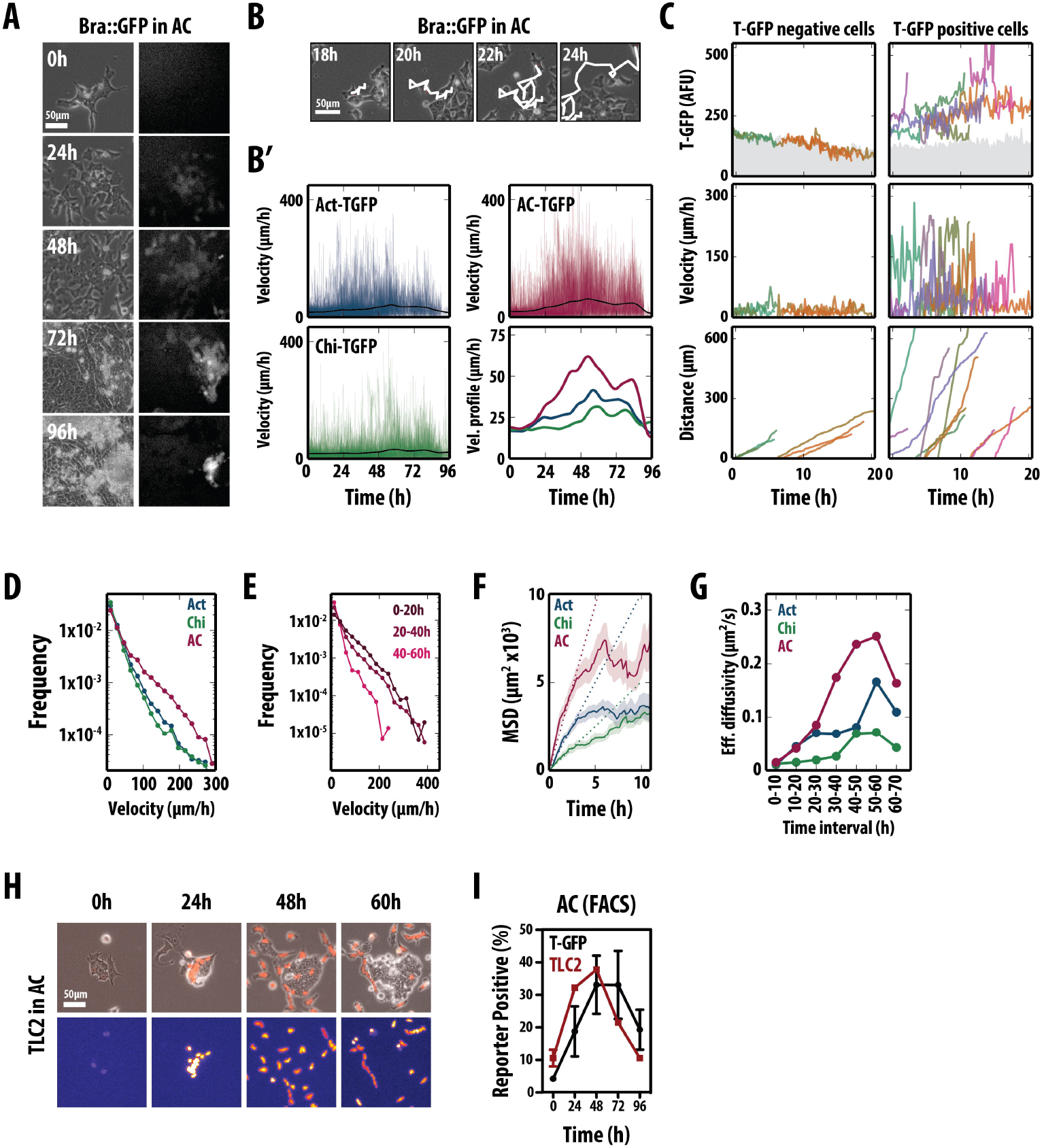
Live microscopy of Bra::GFP and a β-Catenin transcriptional reporter (TLC2) following mesendodermal differentiation. (A) Still images from live imaging of Bra::GFP mESCs in Act/Chi conditions (movie M1) showing phase contrast (left column) and fluorescence (right column). Cells were manually tracked (B) and their velocities (B’, C middle panel), GFP expression (C, top panel) and distance travelled (C, bottom panel) were measured. The average velocities for all tracked cells in Act (blue), Chi (green) and Act and Chi (red) (B’, bottom right) show that cells in Act and Chi have on average the highest peak velocities which are reached earlier. In C, each cell is represented by a colour and it is possible to see that all cells move, but only cells that express Bra move with a high velocity; there seems to be a relationship between the levels of Bra expression and the velocity and distance travelled by individual cells. (D) Distribution of velocities of individual cells under different conditions. Cell in Act and Chi have a higher proportion of fast movers. The velocity of the cells increases with time (E) (F) Mean-squared displacement (MSD) curves representing the range of movement of individual cells, when considering all cells tracked over the entire course of the experiment. (G) Effective diffusion coefficients at different time intervals. The coefficients were estimated by fitting straight lines to the MSD curves obtained when considering cells at different 10h intervals. (H) Live-imaging of the TLC2 Wnt/ß-Catenin reporter in Act/Chi (Movie M2). As cells are differentiated, they begin to up-regulate the reporter before the cells initiate the EMT. The reporter is then down-regulated as cells leave the colonies. (I) FACS analysis of both the Bra::GFP and TLC2 reporter revealing the activation of β-Catenin transcriptional activity relative to the activation of Bra within a population of cells. Average of three replicate experiments ± SD. Scale-bar in all images represents 50 µm.

After 24-48h of Act/Chi treatment, a large proportion of cells undergo EMT-like movements and become openly motile (Fig. 4A; Movie M1). In these conditions, up-regulation of the Bra::GFP reporter was heterogeneous and occurred in cells that were in the process of or undergoing an EMT event within a temporally restricted window (Fig. 4A). The time to the acquisition of a GFP-positive state was fixed, always spanning the second to early third day of differentiation. This correlation between Bra expression, EMT and movement is repeated when cells are treated with Chi or Act individually (Fig. S4A). In Act conditions however, fewer cells up-regulated the Bra::GFP reporter (see Fig. 1A) and the EMT-like events occurred between 48 and 72h after initiation of treatment (Fig. S4A). Up-regulation of the reporter was on average more rapid in Act/Chi conditions than with either Chi or Act alone.

To quantify the characteristics of cell movement and GFP expression, individual cells were manually tracked for long periods of time at a high temporal resolution (∼6 frames/hour, Fig. 4B) and their Bra::GFP levels, instant velocities (the velocity of a cell from frame to frame) and motility (or diffusivity) were measured (Fig. 4B’-G & Supplemental Fig. S4B,C). Population-wise, cells in Act/Chi achieved higher instant velocities much earlier and sustained them for a larger period of time than in either Act or Chi alone (Fig 4B’,D). Binning the velocities based on the time at which the cell analysis began (0-20h, 20-40h, 40-60h) revealed that cells did not have high velocities from the very beginning but these built up as time progressed (Fig 4B’,E).

Although the ability of Act or Chi to alter the instant velocities of individual cells was clear in our analysis, we resorted to statistical methods of random motion particle tracking to provide an objective measurement of the differences in cell motility between the different stimulation conditions (Fig. 4D-G). In particular, we computed the mean-squared displacement (MSD) curves for each cell (Fig. 4F), which, despite being less intuitive than cell velocities, provide a robust analytical framework for rapidly moving cells. When considering all cells tracked over the entire course of the experiment, the evolution of the MSD in Act/Chi is much higher than in the case of individual Act or Chi stimulation (Fig 4E) i.e. cells treated with Act/Chi travel, on average, much larger distances in the same amount of time than cells in either Act or Chi. In addition, in the three cases (Act, Chi or Act/Chi), the initial MSD increases approximately linearly with time, a sign of diffusive movement, and after the first three hours these plateau (Fig. 4F). This levelling out of the MSD is due to a combination of both cell confinement as well as to the fact that we have only tracked cells that remain within the field of view (and thus we underestimate the MSD over long periods of time). Therefore, for each condition, we estimated an effective diffusivity (Supplemental Fig. S4). The evolution of the coefficient of diffusion over time indicates that cells in Act/Chi acquire a much larger degree of motility than cells in Act or Chi and that they do so both much earlier and for a longer period of time (Fig. 4G).

In addition, we studied the relationship between Bra expression and the dynamic behaviour of individual cells treated with Act/Chi. We found a correlation between Bra expression and cell movement and also that those cells with higher velocities had higher levels of the Bra::GFP (Fig. 4C). Consistently, individual cells with levels of Bra::GFP above the average covered much larger ground (Fig. 4C, lower panel).

We further analysed the trajectories of the cells to understand whether there was a particular bias to cell motion in terms of directionality and persistence (a continued direction of movement on a cell by cell basis; Fig. 4H & S4D). In all conditions tested (including both an N2B27 and Serum LIF control), the distribution of turning angles was far from isotropic (Fig. S4D, all conditions had a *p-value* < 0.00001 for the Kolmogorov-Smirnov test against the uniform distribution), a clear indication that regardless of the medium condition, when cells move, they do so with persistence and that the main difference between different conditions is the velocity and the ground covered by individual cells (Fig. S4D).

The importance of Wnt/β-Catenin signalling in the onset of Bra expression (Arnold et al., 2000; Yamaguchi et al., 1999) (and here) led us to monitor Wnt/β-Catenin signalling during differentiation using an H2B-TCF/LEF-mCherry reporter over time (TLC2; Fig. 4H,I) (Faunes et al., 2013; Ferrer-Vaquer et al., 2010). Cells initially expressed low, heterogeneous levels of fluorescence. Over time however, reporter expression increased within the centre of colonies until cells began to disperse from the edges as they initiated EMT (Fig. 4H & Movie M2). The activity of the Wnt-reporter in the transitioning cells was down-regulated following their exit from the colony, correlating with Bra::GFP down regulation in highly motile cells (Fig. 4I).

Our observations and analysis provide information on the temporal order of events with respect to Bra induction, β-Catenin signalling and the EMT: a) cells increase the ß-Catenin transcriptional activity, b) cells express Bra and initiate the EMT event and c) cells reduce transcriptional activity of ß-Catenin and Bra as cells migrate away from the colony and become motile.

### Nodal/Activin and Wnt/β-Catenin signalling provide a link between the EMT and Bra expression

Our observations reveal a close relationship between the onset of Bra expression, the associated EMT and the activation of Wnt signalling. To establish a functional relationship between these three events, we first used CyclosporineA (CsA) to inhibit the EMT during the differentiation process and assessed the expression of Bra::GFP by FACS (Fig. 5A). CsA has been shown to inhibit Calcineurin thereby preventing both the phosphorylation and therefore the activation of nuclear factor of activated T-Cells (NFAT)(Clipstone and Crabtree, 1992; Li et al., 2011) and the transition from an epithelial towards a mesenchymal state (Mancini and Toker, 2009). In our experiments, CsA delayed the onset of Bra::GFP expression induced by Act alone and reduced the number of GFP-positive cells to a maximum of <10% at the end of the treatment (Fig. 5Aa). In the case of Chi, CsA resulted in an immediate reduction of Bra::GFP expression throughout the whole period of observation to levels similar to those observed with Act and CsA (Fig. 5Ab). Although simultaneous Act/Chi and CsA treatment showed an initial decrease in the proportion of GFP-positive cells after 24 hours, CsA treatment appeared only to delay the onset of Bra::GFP induction and shift the GFP-positive distribution by 48 hours (Fig. 5Ac).

**Figure 5:**
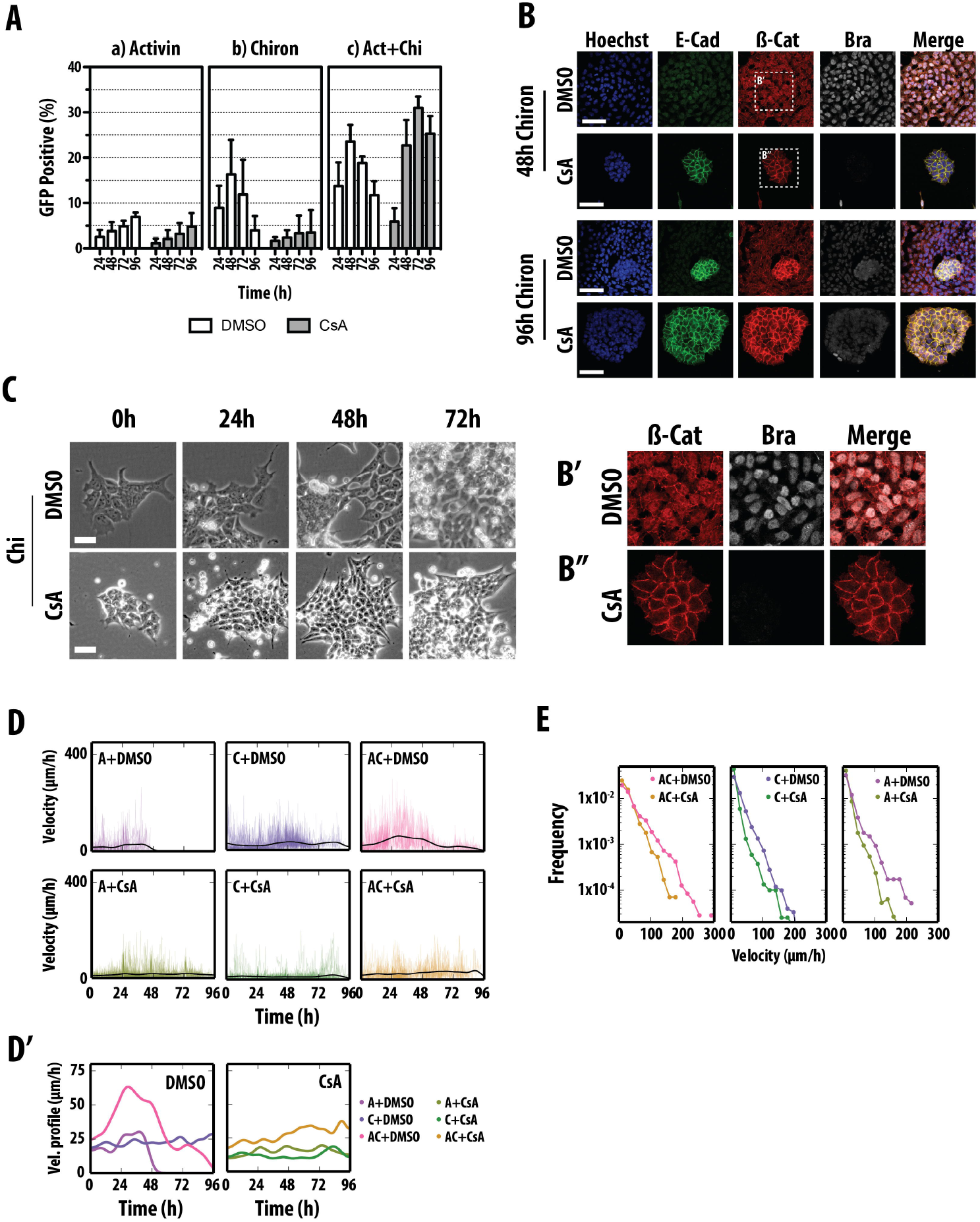
An EMT event is required for Bra::GFP expression. (A) Expression of Bra::GFP in mESCs subject to (a) Act, (b) Chi and (c) Act/Chi, in the presence of CsA (3 μM; grey) or vehicle (DMSO; white) analysed by FACS; s.d shown, n = 3. (B,B’,B’’) E14Tg2A mESCs subject to Chi in the presence of CsA (3 μM) or DMSO, stained for E-Cadherin, β-Catenin, Bra and Hoechst; scale = 50 μm. In the presence of CsA E-Cadherin is not effectively cleared from the membrane, ß-Catenin does not enter the nucleus and there is no effective expression of Bra (B, B’,B’’). (C) Stills from live imaging of E14-Tg2A mESCs in Chi with DMSO or CsA. Notice that in CsA cells stretch out filopodia but do not undergo an EMT; scale = 50 μm. (D) Cell velocities (μm/h) measured from multiple films as in C. Analysis of the average instant velocities shows that CsA reduces the movement of the cells. This is shown in E, as the distribution of instant velocities.

In the presence of Chi, E14-Tg2A cells exhibit a loss of E-Cadherin from the membrane and nuclear relocalisation of β-Catenin coincident with the EMT and the onset of Bra expression (Fig. 5B,B’ & Fig. S2A). Instead, cells in both Chi and CsA did not lose E-Cadherin expression and for the most part, β-Catenin remained membrane-associated (Fig. 5B,B’’). Cells in these conditions display low expression of Bra and of Bra::GFP (Fig. 5A,B,B’’) and show that a release of β-Catenin from the adherens junctions is a prerequisite for its effect on Bra expression. On the other hand, cells treated with Act and CsA showed limited up-regulation of Bra (the same as we observe with Bra::GFP) with incomplete degradation of E-Cadherin (Fig. S5A). However, CsA had a much lesser effect in the presence of Act/Chi resulting in similar, though delayed, E-Cadherin and membrane β-Catenin degradation to the DMSO control, and expression of Bra (Fig 5A & Fig. S5B). The combined effect of Act/Chi on Bra expression although unintuitive can be interpreted as following: Act loosens the adherens junctions, that release a small amount of β-Catenin from the membrane, and Chi amplifies this effect. In the presence of Chi and CsA, there is no β-Catenin released because this requires the action of Act: in Act and CsA but no Chi, there is no feedback as β-Catenin remains in the membrane and; finally, in the presence of both, Act/Chi, the CsA simply delays the process.

Live imaging of cells under these conditions revealed that although cells treated with either Act or Chi in the presence of CsA begin to form membrane protrusions such as Lamellipodial and Fillopodial projections, suggestive of EMT initiation ((Yilmaz and Christofori, 2009) and as confirmed with immunofluorescence), they remained within tight colonies and did not become motile (Fig. 5B,C). This is supported by the quantitative analysis of the cell movements over time (velocities; Fig. 5D-E and Fig. S5C) which revealed that cells treated with CsA are unable to show the rapid cell motion observed in the DMSO controls, even in the presence of Act/Chi, and were more likely to move under 36 μm/h (the basal velocity of cells) for prolonged periods of time and were retarded in their ability to engage in large cell-steps under all conditions (Fig. 5D-E).

These results, taken together with the inhibitor studies using SB43 (Fig. 1Aiv), suggest that activation of Bra transcription is dependent on a) the ability of individual cells to execute an EMT program b) the loss of E-Cadherin at the cell surface and c) the concomitant nuclear translocation of β-Catenin. Consistent with this conclusion, β-Catenin^-/-^ mutant cells and cells harbouring a transcriptionally inactive β-Catenin, β-Catenin^ΔC/-^, were unable to express Bra in the presence of Act and Chi (Fig. 6A,B). Time-lapse imaging revealed striking differences between the two β-Catenin mutant cell lines and the wild-type (Fig. 6C). Whereas by 24 hours E14-Tg2A mESCs began to show signs of the initiation of the EMT (dispersing cells with a mesenchymal appearance), cells mutant for β-Catenin remained tightly associated (Fig. 6C). As time progressed, the β-Catenin^-/-^ cells showed a much greater reduction in viability compared to the E14-Tg2A control and became motile (albeit much slower) however they remained within close proximity to the colony from which they were dispersing (Fig. 6B). Manual tracking of these cells revealed that the speed at which cells move, and thereby indirectly the distance covered, over the time course was significantly shorter than the E14-Tg2A control, with a velocity much less than 36 μm/hour for most of the time course (Fig. 6D,E, Fig. S6A). Unlike the E14-Tg2A control, the β-Catenin^ΔC^ cell colonies at the initiation of imaging (90% out of the colonies of cells present at the initiation of imaging) formed tight, spherical colonies not dissimilar to those observed in 2iLIF conditions (Fig. 6C). Due to the high degree of compaction within the cell balls of the β-Catenin^ΔC^ mutant line, single cells could not be followed for periods of time that allowed comparisons of their velocities with the other mutant lines and the E14-Tg2A controls. The appearance of the colonies is probably due to enhanced E-Cadherin mediated adhesion in these cells.

**Figure 6:**
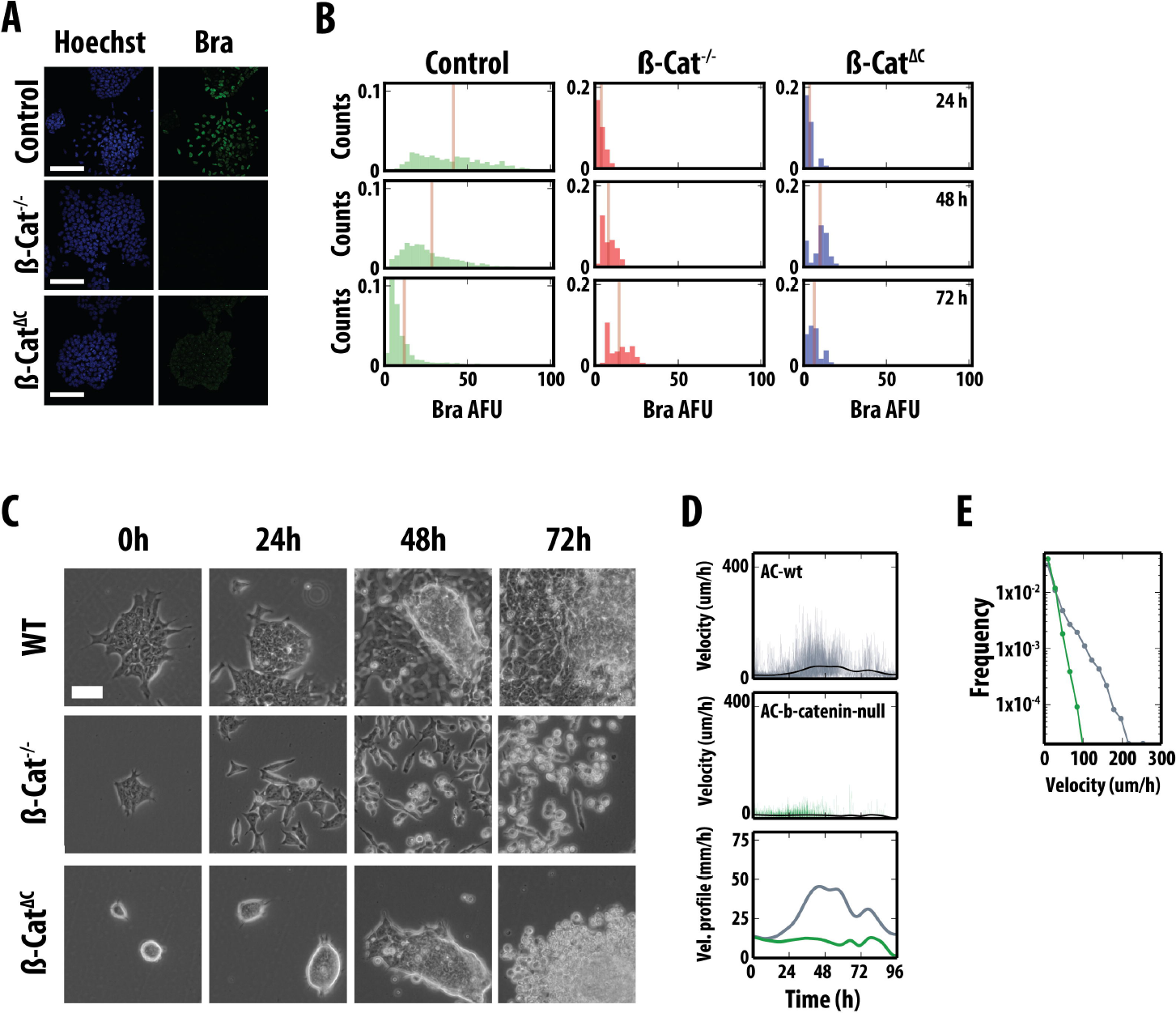
β-Catenin transcriptional activity and expression of Nanog are required for Bra expression and the associated EMT. (A) Bra expression in E14-Tg2A, β-Catenin^-/-^ and β-Catenin^ΔC^ cells differentiated in Act and Chi; Hoechst in the left hand side (B) Time evolution of the distribution of Bra expression in cells as in A. ß-Catenin transcriptional activity is required for Bra expression. See text for details. (C) Live imaging of β-Catenin mutants in Act/Chi. (D) Individual cell velocities (µm/h), and the average velocity profile for each cell line (indicated by colours) in Act/Chi. (E) The distribution of velocities over time. Scale bar = 50μm.

These results, in combination with the chemical genetics approach above, revealed that partial or complete loss of β-Catenin and disruption of its transcriptional activity impairs the ability of ESCs to both upregulate and undergo specification towards a Primitive Streak-like fate.

### Nanog is required for the effect of Bra on the EMT

Our previous analysis on the correlations between Nanog and Bra expression (Fig. 3D,G) and a previous investigation in human ESCs detailing Nanog regulation of Bra (P. Yu et al., 2011) led us to follow the ability of Nanog^-/-^ mESCs to express Bra and undergo EMT by both live-cell imaging and QIA (Fig. S6). QIA analysis shows that following differentiation in Act/Chi, Nanog^-/-^ mESCs express much lower levels of Bra and Sox2 (Fig. S6B). Live imaging of these cells by wide-field microscopy revealed that they undergo an EMT (Fig. S6C), although subtle differences can be observed in the dynamics of cell movements and in the morphology associated with their differentiation (Fig. S6C-E): analysis of the instant velocities revealed that Nanog^-/-^ mESCs were less likely to engage in rapid velocities compared with the wild-type cells although not to the same extent as the ß-Catenin^-/-^ line (Fig. SD,E). These observations suggest that Nanog facilitates the up-regulation of Bra, though its presence is not a requirement for the initiation of the EMT program.

### Brachyury expression is required for rapid cell movement following an EMT

In the embryo, the absence of Bra leads to truncations of the body axis and defects in axial extension, but does not have a major phenotype during gastrulation until the formation of the node (E7.5) (Beddington et al., 1992; Gluecksohn-Schoenheimer, 1938). This led us to test the behaviour of Bra mutant cells in the context of the ES cell differentiation; in particular, we asked whether the EMT program is activated as a consequence or is independent of the initiation of Bra transcription. To test this we analysed the properties of Bra^-/-^ mESCs in culture (Fig. 7).

**Figure 7:**
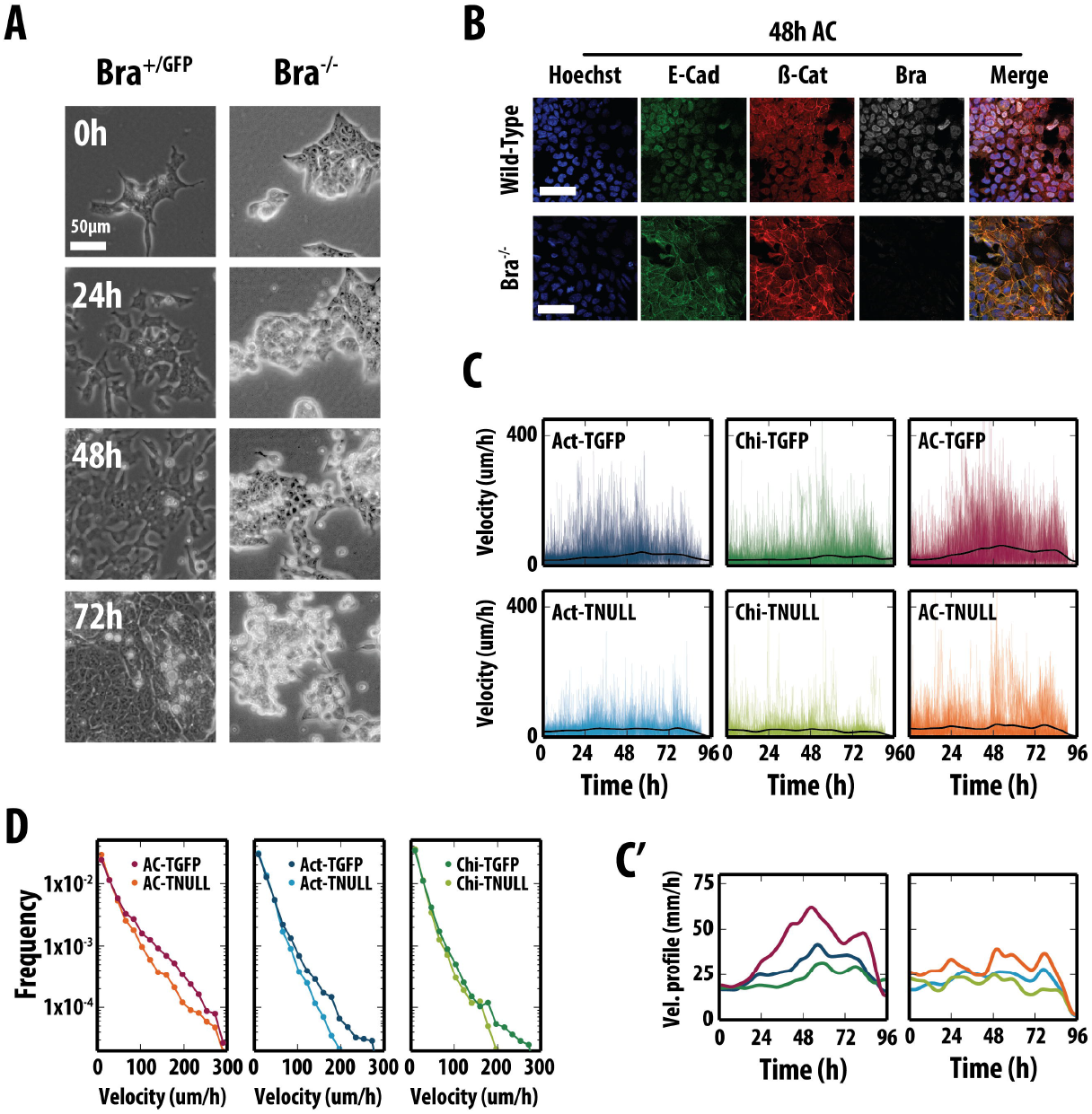
Brachyury is required for EMT-event during mesendodermal differentiation. (A) Phase contrast images from live imaging of Bra::GFP(Bra^+/GFP^) and Bra^-/-^ mESCs in Act and Chi; Bra::GFP stills taken from Fig. 4A. Notice that in the absence of Bra, cells do not undergo a full EMT. (B) E14-Tg2A and Bra^-/-^ mESCs after 48h in Act and Chi, immunostained for E-Cadherin, ß-Catenin, and Bra and imaged by confocal microscopy (Scale bar indicates 50μm, Hoechst marks the nucleus). In the absence of Bra, E-Cadherin and ß-Catenin are not effectively cleared from the membrane. (C, C’) Instant velocities of Bra::GFP cells and Bra null cells in Act, Chi or Act/Chi. The thick black line in the individual graphs in indicates the average velocity for each condition, and is displayed in greater detail in (C’). (D) Histogram of instant cell velocities as measured from frame-to-frame displacements for both the Bra::GFP and Bra^-/-^ mESCs. All the Bra::GFP live imaging data was displayed in Fig. 4.

In all conditions, Bra^-/-^ cells were able to display the characteristics of an EMT, although the proportion of cells undergoing these changes appeared lower than the WT cells and the process was defective. In the presence of Act and Chi, although mutant lines were able to display movements associated with an EMT, immunofluorescent staining revealed that in most cells, the degradation of E-Cadherin and the subsequent nuclear localisation of β-Catenin were impaired (Fig. 7B). Nonetheless, by 96 hours there was a clear decrease of E-Cadherin in the membrane and some β-Catenin above background in the nucleus (Fig. 7B). Manual tracking of the mutant cells over the time-course revealed their average instant velocity in Act and Act/Chi to be slower than the control Bra::GFP cells (Fig. 7C-D & Fig. S7; Bra::GFP cells from Fig. 4) and therefore the distance travelled per time-step was less than the control cells (Fig. 7D). Interestingly, the effect of Chi on the Bra mutant cells was not as pronounced as that seen in Act or Act/Chi conditions: the probability of generating high velocities with Bra mutants was similar to the control cells with Chi treatment (Fig. 7D).

These observations indicate that Bra is not necessary for the EMT, but functions to promote high-velocity motion within Bra expressing cells, possibly facilitating their exit from the Primitive Streak. Together with the inhibitor studies and live-cell imaging of reporter cell lines, these data suggest that Bra, under the control of β-Catenin modulates the instant velocities of the cells as they enter the Primitive Streak fate and may control the expression of a factor, or factors that allow progression of the EMT and therefore specification of the Primitive Streak. It also shows that, for the most part, the effect of Chi on cell movement is not significant and is independent of Bra expression.

## Discussion

It is thought that ESCs provide a unique model system for interrogating developmental processes in culture (Murry and Keller, 2008; Zhu and Huangfu, 2013). However, this possibility rests on the assumption that processes in tissue culture mimic events in the embryo and this has not been yet adequately tested. Here we have applied a combination of live cell microscopy, QIA and FACS to investigate the onset and consequences of Bra expression in differentiating mESCs and analyse to what extent they reflect related events in embryos. In contrast to earlier studies, which have been performed at the level of bulk populations of differentiating cells and mostly focusing on gene expression, we have analysed the dynamics and correlation between gene expression and cell behaviour at the level of single cells. Our the idea is to compare the events in culture with the specification and morphogenetic activity of a related population of cells in the embryo: the Primitive Streak.

In the mouse embryo, the expression of Bra is tightly linked to the process of gastrulation and body extension, in particular, to a population of cells in the epiblast that experience a wave of EMTs associated with directional movement (Lim and Thiery, 2012; Thiery and Sleeman, 2006; Williams et al., 2011). This population sows the precursors for the definitive endoderm and the mesoderm and plays a central role in laying down the axial structure of mammalian embryos (Arnold and Robertson, 2009; J. C. Smith, 2004; Tam and Gad, 2004; Wilson and Beddington, 1997). Throughout gastrulation, Bra is expressed transiently within a moving population of cells, and a similar pulse of expression has been inferred from FACS analysis of cultured ES cells under the influence of Act and β-Catenin signalling (Fehling et al., 2003; Gadue et al., 2006; Hansson et al., 2009). We have confirmed this observation and extended it by showing that the onset of Bra expression is cell autonomous and that, like in the embryo, it is associated with an EMT (Ramkumar and Anderson, 2011; Tam and Gad, 2004; Williams et al., 2011)). Furthermore, we observe in culture that after the EMT, as cells become migratory, they lay down an ECM over which they move. This situation mimics the embryo where cells move over a bed of Fibronectin which has been shown to be important for fate specification (Cheng et al., 2013; Nakaya et al., 2008; Villegas et al., 2013). In the embryo, it is difficult to resolve the source of the Fibronectin, as it could be the epiblast or the ingressing cells. Our results would suggest that it is the ingressing cells that have expressed Bra that make an important contribution laying down the ECM.

In ESCs the expression of Bra is confined to a narrow temporal window, between days 3 and 4. If one assumes that ESCs are in a state similar to that of the preimplantation blastocyst (stage E4.0), the timing of Bra expression in differentiating ES cells, two days later, is similar to that of embryos, between E6.5-7.0, the time at which gastrulation commences (Snow, 1977; Tam and Gad, 2004). There are other features of the process in culture that parallel events in the embryo. In particular the association of Bra expression with β-Catenin signalling (Arnold et al., 2000; Ferrer-Vaquer et al., 2010; Tada and J. C. Smith, 2000), an EMT and the coexpression of both Nanog, and Oct4 with Bra during these processes (Hoffman et al., 2013; Thomson et al., 2011). There are reports that in human ESCs, Nanog is required for Bra expression (P. Yu et al., 2011) and here we have shown that the same is true in mouse ESC where the absence of Nanog dramatically reduces the expression of Bra and affects the behaviour of the cells.

Altogether, our observations support and extend the notion that in vitro differentiation of ESCs into a Bra expressing population, exhibits several parallels with the definition and behaviour of the Primitive Streak during mammalian gastrulation beyond gene expression profiles (Gadue et al., 2006; Izumi et al., 2007). This opens up the possibility of using ESCs to probe the molecular mechanisms linking cell fate and cell behaviour and, by comparing the evolution of the processes in cells and embryos, gain some insights into the emergence of collective behaviour from the activities of single cells.

Our results suggest an interplay between Act and Wnt/ß-Catenin signalling, the EMT and the activity of Bra in the specification and behaviour of cells in the Primitive Streak. Act initiates the EMT and the expression of Bra. The EMT triggers Wnt/ß-Catenin signalling that enhances the effect of Act on Bra which, in turn, promotes cell movement and cell fate (A. L. Evans et al., 2012; Gentsch et al., 2013). This module has the structure of a feed-forward loop. In agreement with these notions, Bra has been shown to control the expression of several components of the cytoskeleton and of canonical/non-canonical Wnt signalling (Hardy et al., 2008; Heisenberg et al., 2000; Shimoda et al., 2012; Tada and J. C. Smith, 2000) which are likely to promote movement and enhance the EMT. Downstream targets of Bra comprise members of the Wnt family which are likely to fuel movement. It is possible that the sluggish movement that we observe in the absence of Bra, is due to the activation of β-Catenin by Chi that might set in motion some of these mechanisms in a Bra-independent manner. In the absence of other elements, also controlled by Bra, the movement is greatly hampered.

### A tissue culture model for Primitive Streak formation?

Differences between the events in the embryo and those in differentiating ESCs can be informative. An example is provided by the long range movement that we observe in differentiating ESCs which is not obvious in the embryo. During gastrulation, after their EMT, cells expressing Bra do not display long range movement as individuals but rather jostle as a group towards the proximal posterior pole and then ingress through the Primitive Streak (Williams et al., 2011). However, when they are explanted and placed onto ECM covered culture dishes the same cells can be observed to move individually, without a preferred direction but with some persistence/diffusivity (Hashimoto et al., 1987) in a manner that is very reminiscent to what we have described here for differentiating ESCs. These observations suggest that a main difference between Bra expressing ESCs and those in the embryo, is the confinement of the latter that restricts their movement and forces them to behave as a coherent collective, rather than becoming dispersed individual cells, as they do in the culture. It is interesting that the average velocity of the differentiating ESCs cells in Act/Chi (maximum average instant velocity of ∼60 µm/h; Fig. 4B’) is within the same order or magnitude as that of the cells from Primitive Streak explants (average of 50 µm/h on extracellular matrix-coated surfaces) (Hashimoto et al., 1987) and of migrating mesodermal cells within the embryo (46 µm/h)(Nakatsuji et al., 1986). Although it is important to note that in our experiments, we were able to see a small proportion of cells that were able to travel at ∼400 µm/s, albeit for short durations of time (Fig. 4B’).

We observe a correlation between the levels of Bra and the velocity of the cell. Bra mutant cells are very delayed in migrating, only a few do migrate and when they do, they exhibit lower velocities relative to wild type. Similar observations have been made for cells from Bra mutant primitive streaks (Hashimoto et al., 1987) and indicate that an important function of Bra is to control the movement of the cells. On the other hand the combination of Act and Chiron promotes very high velocities which, in the confinement of the embryo, can result in strong directional forces.

These observations emphasize the importance of confinement in the behaviour of the cells in the Primitive Streak. At the onset of gastrulation the epiblast is a highly packed and dividing cell population. At this stage, movement towards and through the streak is likely to be due to large-scale, spatially constrained tectonic movements of the epiblast as a whole, with mechanical differences between the prospective anterior and posterior parts being responsible for the directional movement of the bulk population that has undergone EMT. Indeed imaging of the process of gastrulation in mouse embryos suggests that once cells have undergone EMT, Bra expressing cells are passively pushed towards the streak in a process that appears to be passive and not require convergence and extension, as it appears to be the case in frogs and fish (Williams et al., 2011). This notion of morphogenetic events underpinned by long range mechanical coordination of cells within tissues has been demonstrated during the convergence and extension movements of the neuroectoderm in the development of the nervous system in the zebra fish (Blanchard et al., 2009). Such long range effects might provide an explanation for the lack of a clear phenotype of Bra mutant cells during gastrulation. While this is likely to be the result of partial redundancy with Eomesodermin (Arnold et al., 2008; Slagle et al., 2011; Teo et al., 2011), it might be also be a reflection that at these early stages, the defects of Bra mutant cells are compensated by, mechanical large tissue coordination due to confinement. Bra mutant cells still undergo an EMT and therefore could be subject to these movements. As we have shown here, Bra mutant cells can initiate an EMT and therefore could be the subject of strong large scale forces that are likely to exist throughout the epiblast. After node formation, however, the strength of these forces might subside and regressing cells, particularly during axis extension, may come to rely more on the propulsion and navigational abilities driven by Bra. This is supported by the behaviour of mosaics of cells with different levels of Bra in WT embryos (Beddington et al., 1992; Wilson et al., 1995; Wilson and Beddington, 1997) and by our observation of a correlation between levels of Bra expression and cell velocity. Also, in agreement with this, wild type cells are outcompeted by cells expressing higher levels of Bra in the early gastrula (Wilson and Beddington, 1997). Therefore, cells with higher motility might be more prone to escape the tectonic movements of the tissue can ‘overtake’ WT cells.

Our results highlight the experimental possibilities provided by differentiating ESC as a first approximation to understand the mechanisms underlying events that, like gastrulation, are difficult to access in the embryo. Having established the similarities between the two systems, it will be important to exploit them to see how one could reproduce the Primitive Streak in culture through, for example, confining the movement of differentiating ES cells and attempting to create directionality by imposing spatially constrained forces.

## Acknowledgements

We thank F. Bonkhofer for assistance with qRT-PCR, Drs A. K Hadjantonakis, P. Hayward, P.F. Machado, S. Muñoz-Descalzo & C. Schröter and the AMA lab for comments and discussions, Daniel Martinez Gatell for assistance in the manual tracking of cells, Carl Zeiss for technical support, G. Keller for providing the Bra::GFP line, K. Hadjantonakis for the TCF/LEF-mCherry cell line and V. Wilson for the Bra mutant cell line. This research is funded by an ERC advanced grant to AMA.

## Author Contributions

AMA and DAT outlined the project and the experiments; DAT performed the experiments and analysed the data; JPM and PR developed the software for the analysis of the data; DAT, PR, JPM & ED analysed the data. DAT and AMA wrote the manuscript. All authors approved the final manuscript.

## Supplemental Figure Legends

**Fig. S1: Cells cultured in N2B27 are unable to express the Bra-GFP reporter.** Bra::GFP mESCs were plated and differentiated for 48 hours in either N2B27 supplemented with Act/Chi (positive control; Blue line left panel) or in N2B27 (Red line, right panel) without any other factors. The percentage of GFP-positive cells for Act/Chi or N2B27 conditions are shown. An E14-Tg2A wild-type control is displayed in each panel (tinted grey profile).

**Fig. S2: The status of Brachyury, E-Cadherin, Fibronectin and F-Actin in pluripotent and differentiating E14-Tg2A mESCs.** (A) E14-Tg2A mESCs in LIF BMP or differentiated in Act and Chi for 48 hours, stained for Bra (Green) and E-Cadherin (E-Cad; Red) and imaged by confocal microscopy. Act and Chi treatment alters results in Bra expression and a loss of E-Cad from the membrane. (B) E14-Tg2A mESCs cultured in serum and LIF, stained for Fibronectin (green) and with Phalloidin to mark F-actin (red). The pluripotent state is characterized by low Fibronectin and F-actin localized to cell-cell boundaries, lamellipodia and protuding fillopodia; see magnified region (i). (C) A montage illustrating the z-section (in 1 µm increments) of the colony imaged in Fig. 2C showing all fluorescent channels merged (top left), Hoechst and Fibronectin (top right), Hoechst and Bra (Bottom left) or Hoechst and Phalloidin (Bottom right). Hoechst, Fibronectin, Brachyury and Phalloidin are coloured blue, green, white and red respectively. (D) The colony in Fig 2C magnified showing two 320 µm x 13.75 µm sections through the colony (indicated by yellow hashed lines). See text for details. Hoechst was used to mark the nucleus; scale-bar indicates either 50 µm in A and B, or 100 µm in C and D.

**Fig. S3:** The Pearson Correlation Coefficient between for the correlations between Nanog and Oct4 (top), Nanog and Sox2 (bottom) for the different time points in Act, Chi and Act/Chi. The horizontal line represents the correlation for LIF BMP. Differentiation produces strong correlations between Nanog and Oct4 conditions whereas the correlation between Nanog and Sox2 increases with time.

**Fig. S4:** (A) Still images from live imaging of Bra::GFP mESCs in Act/Chi conditions (Movie M1) showing phase contrast (Top rows) and fluorescence (Bottom rows). Act treatment results in lower levels of fluorescence compared with Chi or Act/Chi (Fig. 4). (B) The number of cells tracked per condition over time. (C) The Mean Square Displacement (MSD) of individual cells. Cell traces were separated into 10 hour time-intervals based the time of tracking initiation; the number of cells within each time-interval is displayed. The start of each cell trace within each time-interval were aligned. See text for details. (D) The distribution of turning angles for all cells treated with Act, Chi and Act/Chi. A Serum-LIF control from the TLC2 reporter cell line is included for comparison. There appears to be no bias in the direction cells move with respect to the conditions in which they are placed. Scale-bar = 50 µm.

**Fig. S5:** (A,B) E14-Tg2A mESCs treated with Act (A) or Act/Chi (B) in the presence of CsA (3 μM) or DMSO for 48 and 96h and stained for E-Cadherin (E-Cad), ß-Catenin, Bra and Hoechst. (C) E14Tg2A mESCs subject to Act or in the presence of CsA (3 μM) or DMSO, stained for E-Cadherin, β-Catenin, Bra and Hoechst. In the presence of CsA E-Cadherin is not effectively cleared from the membrane, ß-Catenin does not enter the nucleus and there is no effective expression of Bra (B, B’,B’’). (C) Stills from live imaging of E14-Tg2A mESCs in Act with DMSO or CsA. As with Chi (Fig. 5C) cells in CsA stretch out filopodia but do not undergo an EMT; (D) The number of cells tracked in each condition over time. Scale bar in all images = 50 µm.

**Fig. S6:** (A) The number of cells tracked in each condition over time for wild-type, Nanog^-/-^ and ß-Catenin^-/-^ mESCs. (B) Time evolution of the distributions of the expression of Brachyury in Nanog^-/-^ mESCs. Nanog^-/-^ mESCs treated with Act/Chi for 24, 48 and 72 hours were stained for Bra and nuclei segmented based on hoechst staining. The average pixel intensity for each fluorescent channel (arbitrary fluorescence units: AFU) was quantified and the intensity of Bra displayed as histograms for each time-point the bisecting orange lines in each histogram corresponds to the mean fluorescence levels. (C) Live imaging of Nanog mutants in Act/Chi. (D) Individual cell velocities (µm/h), and the average velocity profile for each cell line (indicated by colours) in Act/Chi. (E) The distribution of velocities over time. For comparison, the velocities and distribution of velocities for the Wild-Type and ß-Catenin mutant cell lines (Fig. 6) are included in D and E. Scale bar = 50μm.

**Fig. S7:** (A) The number Bra^-/-^ mESCs tracked from the live-cell imaging (Fig. 7) in each condition over time. (B) The Mean Square Displacement (MSD) of individual cells. Cell traces were separated into 10 hour time-intervals based the time of tracking initiation; the number of cells within each time-interval is displayed. The start of each cell trace within each time-interval were aligned. See text for details.

## Supplemental Movie Legends

**Movie M1: Live imaging of Bra::GFP cells cultured in Act/Chi.** Bra::GFP mESCs were plated in N2B27 supplemented with Act/Chi. As time progresses, cells begin to display EMT-like movements, become openly motile and begin to express Bra::GFP. Time in hours.

**Movie M2: Live imaging of the ß-Catenin transcriptional reporter line (TLC2) following treatment with Act/Chi.** The TCF/LEF::mCherry (TLC2) mESC reporter line was plated in N2B27 supplemented with Act/Chi. Cells initially expressed low and heterogeneous levels of the reporter which increased over time in the cells that would eventually undergo an EMT event and disperse from the colony. Following colony exit, cells began to down-regulate the reporter. Time in hours.

## Supplementary Table and Legend

**Table S1:**
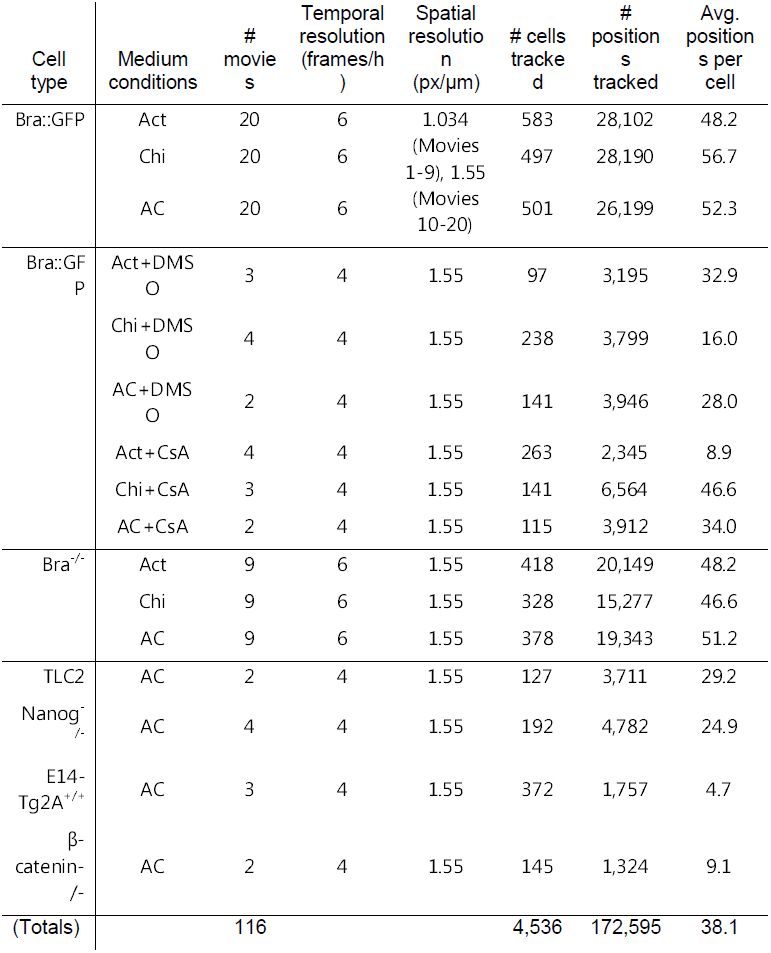
Summary of the tracking data collected. For each cell line used in the live-cell imaging experiments within this investigation (left most column; ‘cell type’), the information relating to the number of movies analysed, the temporal resolution (frames/h), spatial resolution (pixels/µm), the number of cells tracked, positions tracked and the average number of positions per cell for each medium condition (second column from the left; Medium conditions) to which the cells were exposed was tabulated. The total number of movies analysed and the total number of cells tracked throughout the investigation is recorded at the bottom of the table. This table can be used in conjunction with the number of cells tracked over time, displayed in Figs. S4B, S5D, S6A and S7A.

